# Bacteria boost mammalian host NAD metabolism by engaging the deamidated biosynthesis pathway

**DOI:** 10.1101/489674

**Authors:** Igor Shats, Juan Liu, Jason G. Williams, Leesa J. Deterding, Chaemin Lim, Ethan Lee, Wei Fan, Marina Sokolsky, Alexander V. Kabanov, Jason W. Locasale, Xiaoling Li

**Affiliations:** Signal Transduction Laboratory and Epigenetics and Stem Cell Biology Laboratory National Institute of Environmental Health Sciences, Research Triangle Park, NC 27709, USA.; Department of Pharmacology and Cancer Biology, Duke University School of Medicine, Durham, NC 27710, USA.; Mass Spectrometry Research and Support Group, Epigenetics and Stem Cell Biology Laboratory National Institute of Environmental Health Sciences, Research Triangle Park, NC 27709, USA.; Center for Nanotechnology in Drug Delivery, University of North Carolina, Chapel Hill, NC 27514, USA.

**Author notes:** Corresponding authors (I.S.), (X.L.).

## Abstract

Nicotinamide adenine dinucleotide (NAD), a cofactor for hundreds of metabolic reactions in all cell types, plays an essential role in diverse cellular processes including metabolism, DNA repair, and aging ^1^. NAD metabolism is critical to maintain cellular homeostasis in response to the environment, and disruption of this homeostasis is associated with decreased cellular NAD levels in aging ^2^. Conversely, elevated NAD synthesis is required to sustain the increased metabolic rate of cancer cells ^3,4^. Consequently, therapeutic strategies aimed to both upregulate NAD (i.e. NAD-boosting nutriceuticals) or downregulate NAD (inhibitors of key NAD synthesis enzymes) are being actively investigated ^5–10^. However, how this essential metabolic pathway is impacted by the environment remains unclear. Here, we report an unexpected trans-kingdom cooperation between bacteria and mammalian cells wherein bacteria contribute to host NAD biosynthesis. Bacteria confer cancer cells with the resistance to inhibitors of NAMPT, the rate limiting enzyme in the main vertebrate NAD salvage pathway. Mechanistically, a microbial nicotinamidase (PncA) that converts nicotinamide to nicotinic acid, a key precursor in the alternative deamidated NAD salvage pathway, is necessary and sufficient for this protective effect. This bacteria-enabled resistance mechanism that allows the mammalian host to bypass the drug-induced metabolic block represents a novel paradigm in drug resistance. This host-microbe metabolic interaction also enables bacteria to dramatically enhance the NAD-boosting efficiency of nicotinamide supplementation *in vitro* and *in vivo*, demonstrating a crucial role of microbes, gut microbiota in particular, in organismal NAD metabolism.

---

NAD is an essential cofactor in hundreds of redox reactions. It is also consumed by DNA repair enzymes (i.e. PARPs, poly(adenosine diphosphate- ribose) polymerases) and by protein deacylases (e.g. sirtuins) to regulate many fundamental cellular processes, including energy metabolism, genome stability, and circadian rhythms ^1^. Mammalian cells are capable of synthesizing NAD from the amino acid tryptophan (*de novo* pathway) or from nicotinic acid (NA, Preiss-Halder deamidated salvage pathway). However, the main cellular source of NAD is its salvage from nicotinamide (NAM, amidated salvage pathway), where nicotinamide phosphoribosyl transferase (NAMPT) is the rate limiting enzyme ^11^ (Fig. 1A).

**Figure 1.**
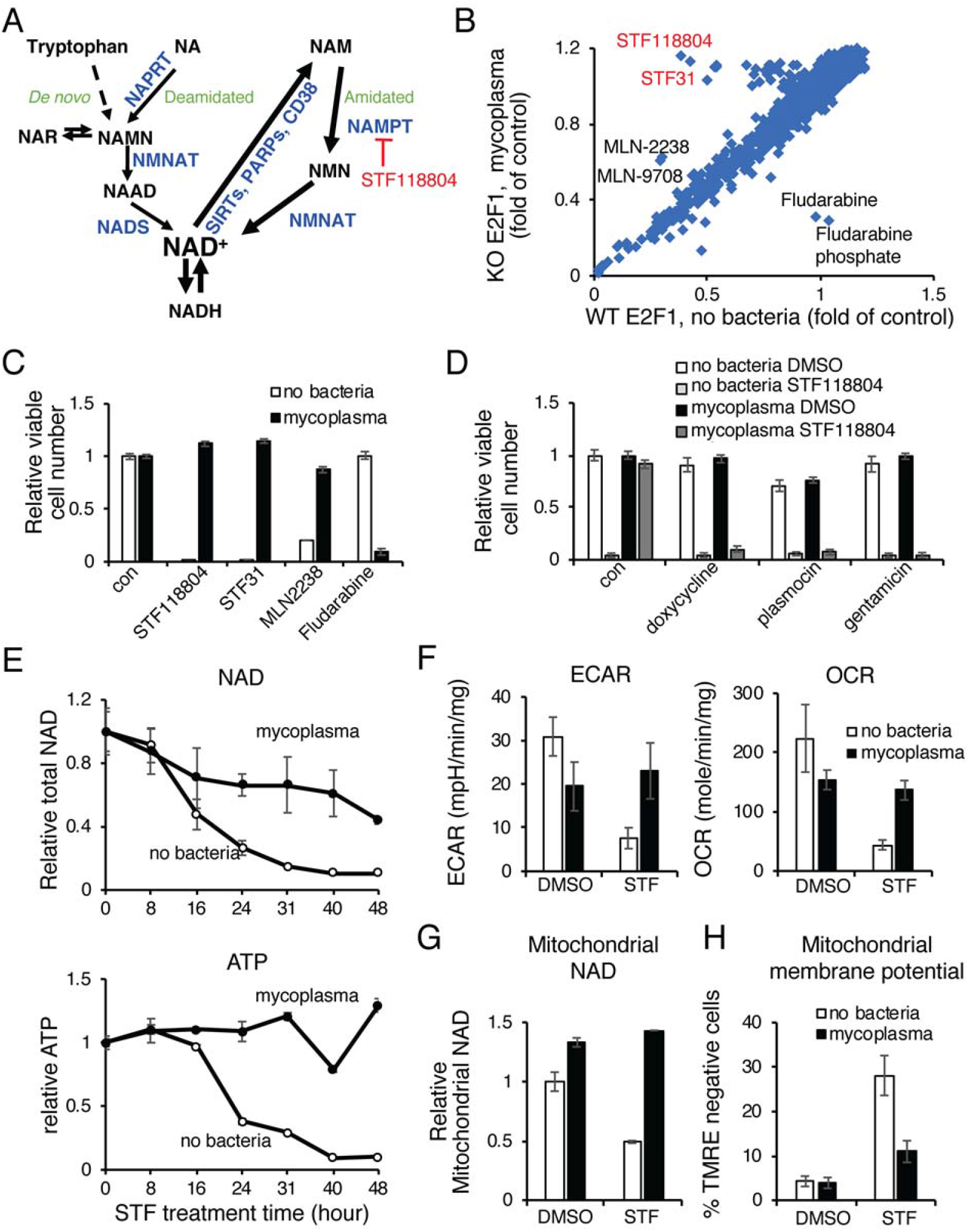
Mycoplasma infection confers host cells with resistance to NAMPT inhibitors by preventing NAD and energy depletion. (A) NAD biosynthesis pathway. NAR: nicotinic acid riboside; NAMN: nicotinic acid mononucleotide; NAAD: nicotinic acid adenine dinucleotide; NADS: NAD Synthetase; NMNAT: Nicotinamide Nucleotide Adenylyltransferase. (B) Drug screen in H1299. WT and E2F1 KO H1299 cells were treated with 1 ŪM compounds from the bioactive compound library and viability was measured 48 hours later by CellTiter-Glo (CTG) assay. E2F1 KO cells were subsequently found to be infected with mycoplasma. (C) Mycoplasma infection confers resistance to NAMPT and proteasome inhibitors but sensitizes to fludarabine. CRC119 cells incubated with a supernatant from a mycoplasma-infected culture or a control medium, and treated with 100 nM STF118804, 1 μM STF31, 50 nM MLN2238 or 1 μM fludarabine or control for 68 hours. (D) Elimination of mycoplasma sensitizes cells to NAMPTi-induced toxicity. Clean and mycoplasma-infected CRC119 cells were treated with 1 μg/ml doxycycline, 25 μg/ml plasmocin, or 400 μg/ml gentamicin for 24 hours, then co-treated with antibiotics and 100 nM STF118804 for an additional 48 hours. (E) Mycoplasma prevent NAMPTi-induced NAD and ATP depletion. Total cellular NAD (NADH + NAD+) and ATP levels were measured in uninfected (no bacteria) or mycoplasma-infected CRC119 cells treated with 100 nM STF118804 (STF). (F) Mycoplasma prevent NAMPTi-induced inhibition of glycolysis (ECAR) and oxidative phosphorylation (OCR). Uninfected clean or mycoplasma-infected CRC119 cells were treated with 100 nM STF118804 (STF) for 48 hours. The basal extracellular acidification rate (ECAR) and oxygen consumption rate (OCR) were measured using Seahorse instrument. (G) Mycoplasma prevent NAMPTi-induced mitochondrial NAD depletion. Relative levels of mitochondrial total NAD were measured after 24 hours treatment with 100 nM STF118804 (STF) or DMSO control. (H) Mycoplasma prevent NAMPTi-induced loss of mitochondrial membrane potential. Mitochondrial membrane potential loss (% TMRE-negative cells) was measured by flow cytometry after 24 hours treatment with 100 nM STF118804 (STF) or DMSO control. All data are means and SD of biological triplicates from representative experiments. Each experiment was repeated at least twice.

Cellular NAD homeostasis buffers the constant perturbations from the environment that affect NAD levels. Whereas various cell-autonomous regulatory mechanisms have been described, the impact of microorganisms on mammalian NAD homeostasis remains largely unknown. We now report a mechanism by which mammalian host NAD metabolism is maintained through a host-microbe metabolic interaction that we discovered serendipitously.

We conducted a chemical screen to identify pharmacologic compounds that induce cell death through E2F1, a transcription factor the level of which plays a crucial role in determining cell fate ^12^. As shown in Fig. 1B, in a screen of 2300 bioactive compounds, the top hits that killed wild-type (WT) cells more efficiently than E2F1 knockout (KO) H1299 lung cancer cells were two NAMPT inhibitors, STF31 and STF118804. E2F1 KO cells were also protected from toxicity induced by proteasome inhibitors, MLN-2238 and its pro-drug MLN-9708, but were more sensitive to fludarabine and fludarabine phosphate (Fig. 1B and Table S1). Subsequently, we found that E2F1 KO cells, but not the WT cells, were contaminated with *Mycoplasma hyorhinis*, and the differential responses of WT and E2F1 KO cells to all top hits from our screen were due to mycoplasma contamination rather than E2F1 deficiency (Fig. 1C and Fig. S1). Particularly, in mycoplasma-free CRC119 colon cancer cells, mycoplasma-containing supernatant from an infected culture was sufficient to completely prevent toxicity induced by the two NAMPT inhibitors, STF118804 and STF31, and by the proteasome inhibitor MLN-2238 (Fig. 1C). Mycoplasma also dramatically sensitized the CRC119 cells to fludarabine (Fig. 1C), likely due to the ability of mycoplasma-encoded purine nucleoside phosphorylases to convert fludarabine (a pro-drug) to a highly toxic purine base ^13^. Chronic infection with *M. hyorhinis* purchased from ATCC also protected CRC119 cells from STF118804-induced cell death (Fig. S2) and conferred resistance to three different NAMPT inhibitors (NAMPTi) in two additional cell lines (Fig. S3). Conversely, treatment with structurally diverse antibiotics that eliminate *M. hyorhinis* completely reversed this resistance (Fig. 1D), further confirming that mycoplasma protects host mammalian cells from NAMPTi-induced cell death. Consistent with these observations in cultured cells, *M. hyorhinis*-infected HCT116 xenograft tumors but not the uninfected tumors were protected against STF118804-induced repression of proliferation genes, such as Cyclin A2 (CCNA2) and E2F1 (Fig. S4A). Our drug screen was highly specific and revealed very few strong hits with NAMPT inhibitors being the top hits. This strongly suggested that NAD metabolism is the main cellular pathway affected by mycoplasma infection.

Mycoplasma are common cell culture contaminants and a clinically important component of the human commensal and pathogenic microbiome in various tissues ^14–16^. Mycoplasma presence has also been reported in various cancer types, however, its functional importance remains unclear ^17^. Given the central role of NAD in normal and tumor energy metabolism, the fact that NAMPTi were the top hits in our drug screen, and the importance of this drug class as novel antineoplastic agents, we next sought to elucidate the mechanism underlying mycoplasma-mediated protection from NAMPTi-induced toxicity. We assessed the effects of STF118804 treatment over time and found that mycoplasma attenuated NAD depletion and completely restored ATP levels (Fig. 1E). Bioenergetic profiling demonstrated that this mycoplasma-provided protective effect was accompanied by a rescue of cellular glycolysis and oxygen consumption in STF118804-treated cells (Fig. 1F), suggesting that mycoplasma restore the cellular NAD pool and oxidative phosphorylation. In particular, mycoplasma rescued the STF1188804-induced reduction of NAD in isolated mitochondria (Fig. 1G) and rescued STF118804-induced loss of mitochondrial membrane potential (Fig. 1H). Therefore, mycoplasma infection prevents NAMPTi-induced energy depletion and subsequent cell death by partially rescuing cellular NAD levels. We showed that similar mechanisms are utilized *in vivo*, as NAD depletion in HCT116 xenograft tumors following STF118804 treatment was significantly attenuated in mycoplasma-infected tumors (Fig. S4B).

To further uncover the molecular basis of this mycoplasma-provided protective effect on cellular NAD metabolism, we performed metabolomic analyses of uninfected or *M. hyorhinis*-infected CRC119 cells treated with STF118804 or DMSO vehicle control. Principal component analysis (PCA) revealed that STF118804 induced dramatic metabolic alterations in clean human cells but mycoplasma infection prevented these global changes (Fig. S5A). Remarkably, the top metabolites differentially induced by mycoplasma in cells were two key intermediate metabolites in the Preiss-Handler deamidated NAD biosynthesis pathway, nicotinic acid riboside (NAR) and nicotinic acid adenine dinucleotide (NAAD) (Fig. 2A and 2B). Furthermore, culture medium from mycoplasma-infected cells contained more than 5-fold higher levels of NA and more than 8-fold higher levels of NAR compared to medium from uninfected clean cells (Fig. 2B and S5B), supporting the notion that the increased intracellular levels of NAR and NAAD originated from the increased extracellular NA or NAR. Notably, intracellular STF118804 levels were not affected by mycoplasma (Fig. S5C), indicating that mycoplasma neither degrade STF118804, unlike the reported mechanism of direct drug degradation by bacteria in the case of gemcitabine ^18^, nor inhibit the cellular uptake of this compound.

**Figure 2.**
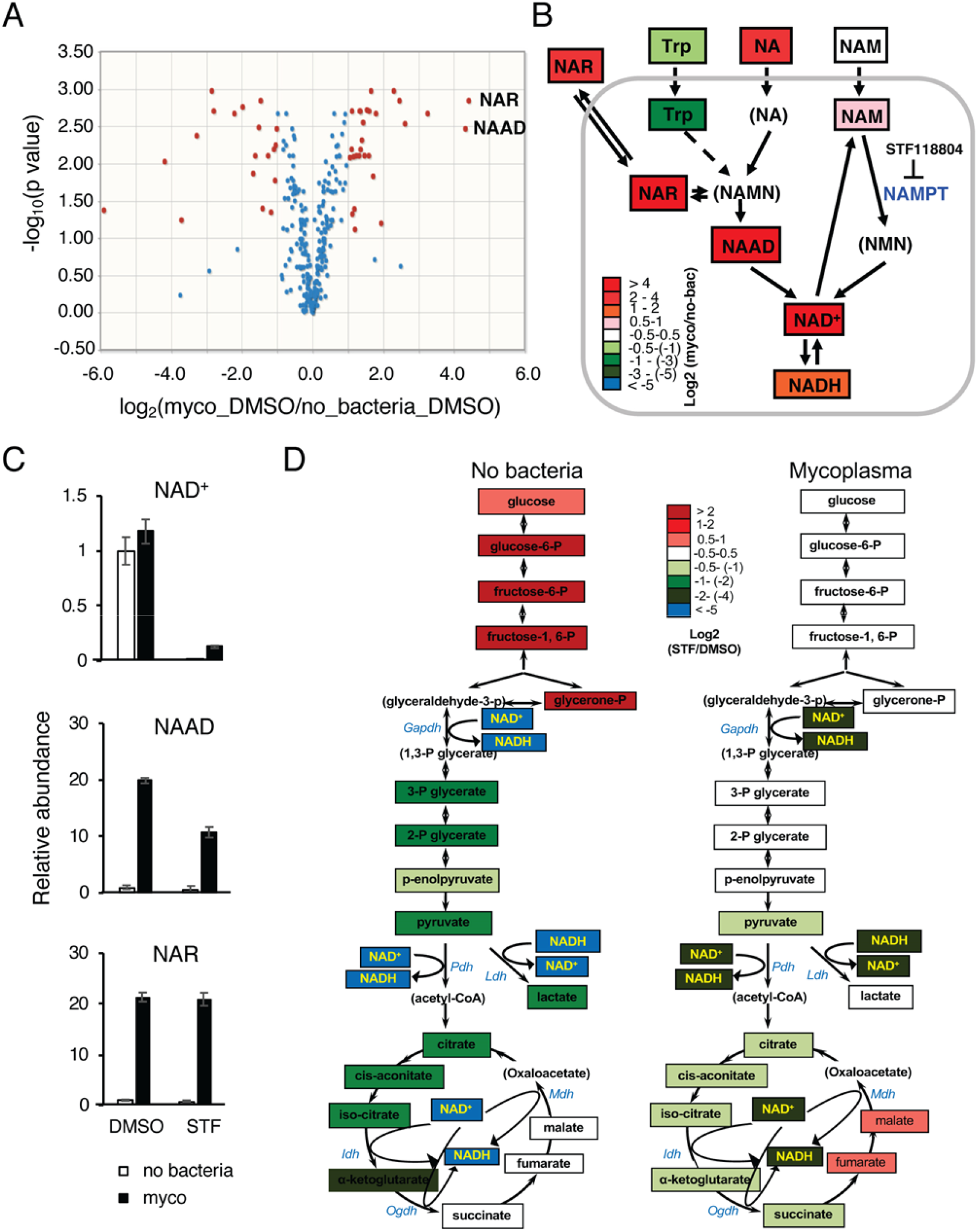
Mycoplasma induce deamidated NAD precursors and prevent NAMPTi-induced NAD and energy depletion. (A) Mycoplasma dramatically increases cellular levels of deamidated NAD precursors. Clean (no bacteria) or M.hyorhinis-infected (myco) CRC119 cells were treated with either 100 nM STF118804 or with DMSO control for 24 hours and relative levels of 330 metabolites were determined by LC-MS. Volcano plot of metabolites from mycoplasma-infected cells (Myco_DMSO) vs. uninfected cells (no bacteria_DMSO). (B) Mycoplasma dramatically increase deamidated NAD precursors inside the cells and in the medium upon STF118804 treatment. The log ratios of the relative abundance of metabolites in clean and infected cells are represented by color scale. Metabolites in parenthesis were not detected in our LC-MS analysis. (C) Relative levels of NAD^+^ and the top differential metabolites (NAAD and NAR). (D) Mycoplasma prevent NAMPTi-induced impairment of energy metabolism. Relative changes in the levels of glycolysis and TCA cycle metabolites following STF18804 treatment are shown for uninfected (left panel) and mycoplasma-infected cells (right panel). The log ratios of the relative abundance of metabolites in indicated pathways in clean and infected cells are represented by color scale. Metabolites in parenthesis were not detected in our LC-MS analysis. See also Table S2 for the complete metabolomics data. All data are means and SD of biological triplicates from a single experiment.

Further analyses unveiled that the mycoplasma-induced increase in the Preiss-Handler deamidated NAD biosynthesis pathway was associated with significant rescue of cellular NAD levels after STF118804 treatment (Fig. 2B and 2C). In particular, STF118804 treatment resulted in a 99.6% drop of cellular NAD levels in clean cells within 24 hours, compared with an 88% drop in mycoplasma-infected cells (Fig. 2C, NAD). We hypothesized that this mycoplasma-mediated partial rescue of cellular levels of NAD (from 0.4% to 12%) is sufficient to sustain the cellular energy metabolism upon treatment with NAMPTi. Consistent with this notion, pathway analysis of differential metabolites in mycoplasma-infected vs. clean STF118804-treated cells identified the TCA cycle as the most significant metabolic pathway rescued by mycoplasma (Fig. S5D). In addition, STF118804 treatment resulted in a complete block of glycolysis at the NAD-dependent GAPDH step in uninfected cells (Fig. 2D, left). Importantly, the STF118804-induced block of glycolysis was rescued and the changes in the TCA cycle were attenuated in mycoplasma-infected cells (Fig. 2D, right). Taken together, our results indicate that mycoplasma primarily affect NAD-mediated energy metabolism in host cells.

The amidated (via NAM) and deamidated (via NA) salvage pathways of NAD biosynthesis are isolated in vertebrate cells due to lack of a nicotinamidase activity that converts NAM to NA (Fig. 1A). In contrast, multiple bacteria species encode nicotinamidases ^19^. Given the dramatic upregulation of the deamidated NAD precursors (NA and NAR) in mycoplasma-infected medium and cells (Fig. S5B and 2B), we hypothesized that protection from NAMPTi by mycoplasma (and possibly other bacteria) may be mediated by bacterial nicotinamidase PncA, which can bypass the NAMPT block by diverting NAM to the deamidated route of NAD biosynthesis. In support of this hypothesis, we found that *E. coli*, like mycoplasma, protected CRC119 cells from toxicity of two different NAMPT inhibitors, STF118804 and FK866 (Fig. 3A). This protection was also accompanied by partial rescue of STF118804-induced NAD depletion in both the cytosol and mitochondria (Fig. S6A) and restoration of both glycolysis and oxidative phosphorylation (Fig. S6B). Importantly, deletion of the *E. coli pncA* gene completely abolished this rescue effect (Fig. 3B). Rescue of viability of NAMPTi-treated cells could also be achieved by addition of conditioned medium (CM) from WT *E. coli* incubated with NAM but not by CM from WT *E. coli* incubated without NAM (Fig. 3C). We detected robust conversion of NAM to NA by *E. coli* in the medium by LC-MS (Fig. S7). Direct supplementation of NA, the product of the PncA enzyme, into culture medium also rescued clean cells from STF118804 toxicity (Fig. 3D). Furthermore, overexpression of *E. coli* PncA in CRC119 cells was sufficient to completely rescue them from STF118804-induced death (Fig. S6C and 3E). In contrast, blocking the deamidated NAD biosynthesis by a NAPRT inhibitor, 2-hydroxynicotinic acid (HNA), abolished the protection from NAMPTi mediated by both *E. coli* and *M. hyorhinis* (Fig. 3F).

**Figure 3.**
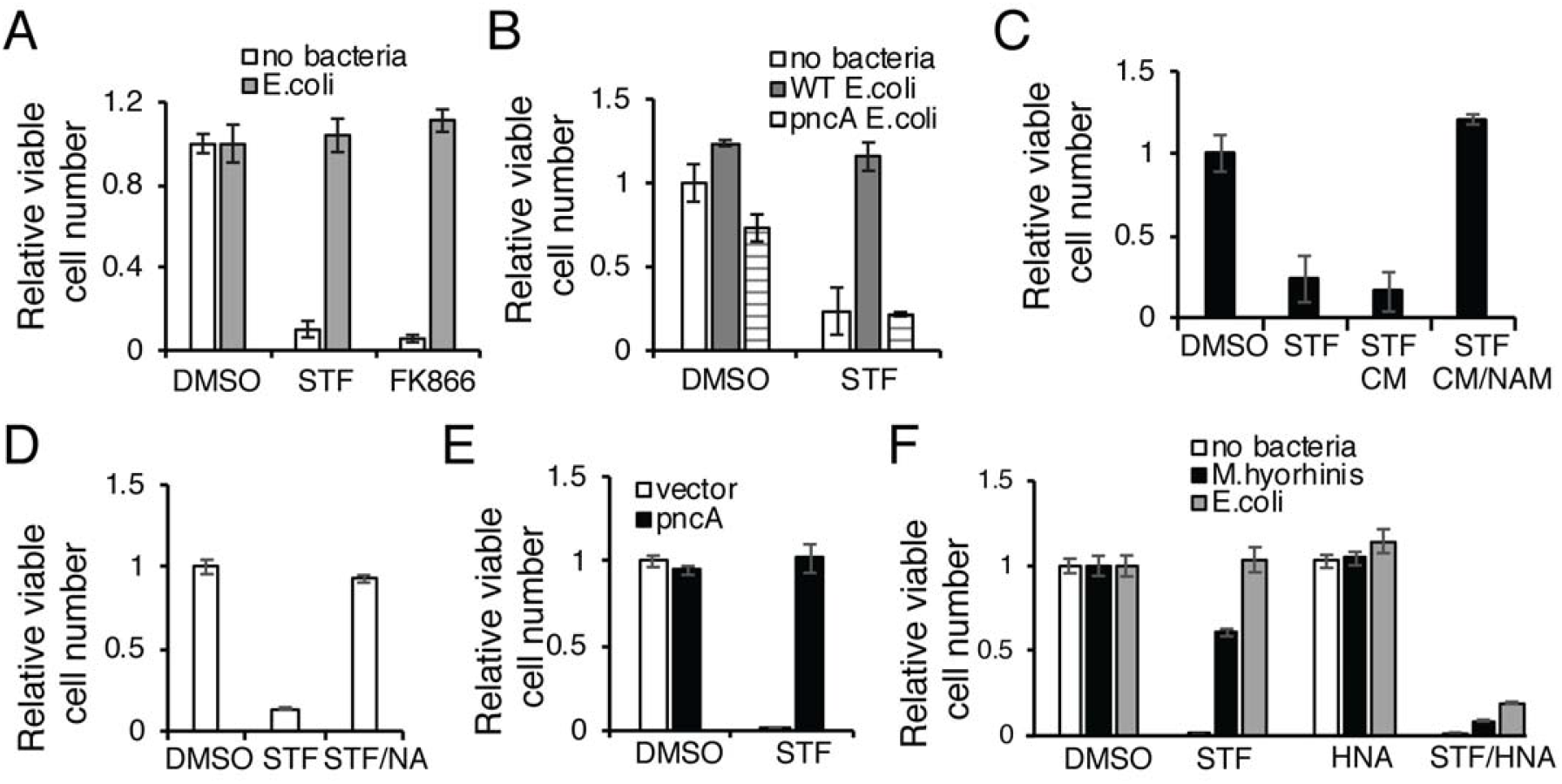
Bacteria rescue NAMPTi-induced toxicity through nicotinamidase PncA. (A) *E. coli* protect cells from NAMPTi-induced toxicity. CRC119 cells were cultured with or without E. coli in the presence of 1 μg/ml gentamycin to prevent bacterial overgrowth. Cells were treated with 100 nM STF118804, 50 nM FK866, or with DMSO control for 42 hours. (B) *E. coli* protect cells from NAMPTi-induced toxicity through PncA. CRC119 cells were treated with 100 nM STF118804 or DMSO control for 48 hours, then treated with control medium, live WT *E.coli*, or pncA KO *E. coli* for additional 24 hours. (C) *E.* coli-provided protection from NAMPTi require NAM. *E. coli* were incubated in EBSS medium with or without nicotinamide (NAM) for 3 hours, then removed by filtering through 0.2 μm filter. The conditioned media (CM) were added for an additional 24 hours to CRC119 cells that were pre-treated for 24 hours with 100 nM STF118804. (D) NA protects cells from NAMPTi-induced toxicity. CRC119 cells were treated with 100 nM STF118804 with or without 100 μM nicotinic acid (NA) for 48 hours. (E) Overexpression of PncA protects cells from NAMPTi-induced toxicity. CRC119 transfected with a control vector or a construct expressing *E. coli* pncA were treated with 100 nM STF118804 (STF) or DMSO control for 72 hours. (F) Blocking the deamidated NAD biosynthesis by HNA abolishes bacteria-provided protection from NAMPTi. CRC119 cells cultured with clean medium, or medium containing *E. coli*. or *Mycoplasma hyorhinis* were treated with 100 nM STF118804, 1 mM of HNA or their combination for 66 hours. Cell viability in (A-F) was measured by CTG assay. All data are means and SD of biological triplicates from representative experiments. Each experiment was repeated at least twice.

Finally, supplementation of bacteria-containing culture medium with isotopically labeled NAM resulted in a robust incorporation of the label into intracellular NA, NAR, NAAD and NAD even when NAMPT was inhibited by STF118804 (Fig. 4A and S6D), demonstrating that *E. coli* and *M. hyorhinis* are capable of converting NAM to NA and promoting the deamidated route of NAD biosynthesis in host cells. In line with this *in vitro* observation, isotopically labeled NAM was rapidly converted to NA and NAAD in the colons when gavaged into microbiota-proficient regular mice (Fig. 4B and 4C, Reg NAM), but this conversion was severely blunted in antibiotic-treated microbiota-depleted mice (Fig. 4C, Abx NAM, Fig. 4D, and Fig. S8). Moreover, in agreement with the previous studies ^6^, NAM supplementation led to more than 100-fold increase in the liver NAAD and a 2.5-fold increase in liver NAD in conventional mice. However, this accumulation of NAAD was abolished and NAD boost was severely attenuated in the livers of antibiotic-treated mice (Figs. 4C and 4D). These results suggest a model in which gut bacteria deamidate dietary NAM to NA, which is then converted to NAAD in gut cells. NAAD is further transported to the liver, where it significantly contributes to hepatic NAD biosynthesis (Fig. 4D).

**Figure 4.**
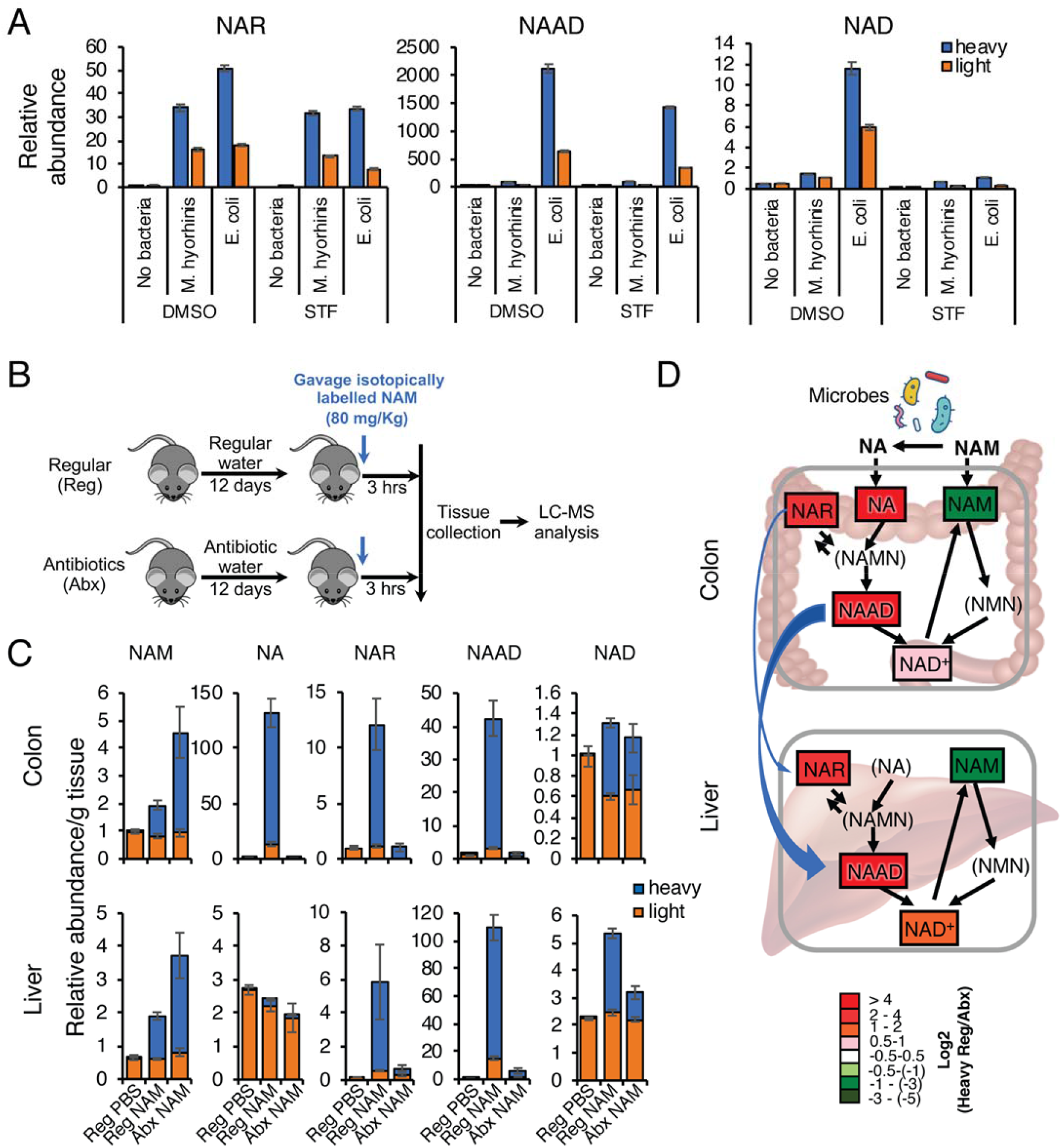
Bacteria boost incorporation of NAM into metabolites in the deamidated NAD salvage pathway and NAD *in vitro* and *in vivo*. (A) Bacteria augment incorporation of NAM into metabolites in the deamidated NAD salvage pathway and NAD. CRC119 cells were infected with the indicated bacteria and treated with 200 nM STF118804 or DMSO control for 24 hours in the presence of 5 mg/l NAM labeled with four deuterium atoms on the pyridine ring. The relative abundance of the indicated unlabeled (light) and labeled (heavy) metabolites was measured by LC-MS. (B-D) Gut microbiota are critical to incorporate dietary NAM into metabolites in the deamidated NAD salvage pathway and NAD *in vivo*. (B) Schematic of the experiment. C57BL/6J mice were treated with either regular water (Reg) or antibiotics-containing water (Abx) for 12 days to deplete gut microbiota. They were then gavaged with 80 mg/kg of isotopically labeled NAM or with PBS control, and dissected three hours later. (C) Relative abundance of unlabeled (light) and labeled (heavy) NAD pathway metabolites in colons and livers were measured by LC-MS (n=5-6 mice/group). (D) Gut microbiota systemically boost the flux of dietary NAM into NAD through the deamidated salvage pathway *in vivo*. The relative abundance of isotopically labeled specific metabolite in colons and livers of regular (Reg) and antibiotics-treated (Abx) mice is normalized to the total level of this metabolite in colons of regular mice. The log ratios of the relative abundance are represented by color scale. Metabolites in parenthesis were not detected in the LC-MS analysis. Data are means and SD from a single in-vivo experiment with 5-6 mice/group.

Collectively, our results demonstrate that bacteria enhance host mammalian cell NAD metabolism by engaging the deamidated route of NAD biosynthesis *in vitro* and *in vivo*, and that this metabolic crosstalk is one of the major interactions between host cells and bacteria. Given the recently reported presence of intratumor bacteria, including mycoplasma, in multiple tumor types ^14,18^, bacteria-mediated resistance described here may have contributed to the failure of NAMPT inhibitors in cancer clinical trials. This finding suggests co-treatment of NAMPT inhibitors with antibiotics as a potential novel therapeutic strategy. Our finding further suggests that development of specific PncA inhibitors for concomitant use with NAMPT inhibitors may provide an even more specific approach to avoid complete microbiome depletion and other undesired side effects of antibiotics. Finally, as commonly used NAD-increasing nutriceuticals nicotinamide mononucleotide and nicotinamide riboside are quickly degraded to nicotinamide after administration ^20^, bacteria-mediated dramatic facilitation of the NAD-boosting activity of NAM supplementation (Fig. 4) demonstrates a crucial role for microbes, particularly gut microbiota, in mediating the efficiency of NAD-increasing nutriceuticals.

## Supporting information

## Acknowledgments

We thank Drs. Traci Hall, Michael Fessler, and Paul Wade, and members of the Li laboratory for critical reading of the manuscript; We also thank Dr. So Young Kim from Duke University Genetic and Chemical Screening Services for construction of E2F1 targeting CRISPR construct and help with the drug screen, and Mr. Michael Johnston, Dr. David Kurtz and the Quality Assurance Laboratory of National Institute of Environmental Health Sciences for detecting mycoplasma contamination in our cultured cells.

## Funding

This research was supported by the Intramural Research Program of National Institute of Environmental Health Sciences of the NIH to XL (Z01 ES102205).

## Authors contributions

IS and XL designed the study, analyzed the results and wrote the manuscript. IS performed all biological experiments with help from EL and WF. JL and JWL performed metabolomic analysis. JGW and LJD performed targeted analysis of NAD pathway metabolites. CL, MS, and AVK developed the formulation of STF1188804 for the *in vivo* studies. All authors critically reviewed the manuscript.

## Competing interests

Authors declare no competing interests.

## Methods

### Cell culture, infections and chemicals

H1299 and 293T cells were obtained from Duke University cell culture facility. HCT116 cells were from ATCC. CRC119 and CRC240 cells were a gift from Dr. David Hsu (Duke University). All cells were grown in RPMI medium with 10% fetal bovine serum supplemented with penicillin and streptomycin. For chronic mycoplasma infection, cells were infected once with *Mycoplasma Hyorhinis* (ATCC^®^ 17981™) and then passaged as usual. For *E.coli* experiments, penicillin and streptomycin in growth medium were substituted by 1 μg/ml gentamycin as we found this concentration to be bacteriostatic. Cells were infected with 1:1000 dilution of overnight stationary phase K12 *E.coli* culture (E. coli Keio Knockout Parent Strain BW25113, #OEC5042 or pncA KO #OEC4987-200827138, Dharmacon). Separate tissue culture reagents bottles and incubator shelves (in addition to secondary containment in larger dishes) were used for the infected cultures. Following the initial discovery of mycoplasma in H1299 E2F1 KO cells, mycoplasma contamination status of all cultures was monitored by real-time PCR monthly ^21^.

STF118804, STF31, FK866, MLN2238, and fludarabine were purchased from Selleckchem. For the *in-vivo* study, STF118804 was purchased from Medchemexpress.

Plasmocin was purchased from Invivogen. All other chemicals were purchased from Sigma.

### Drug Screen

Drug screen was performed at Duke University Genetic and Chemical Screening Services.

Wildtype or E2F1 knockout H1299 cells were plated using a Matrix WellMate onto 384-well plates that had been stamped using a Labcyte Echo Acoustic Dispenser with the Bioactive compound library (Selleckchem), for a final concentration of 1.25 μM in duplicate plates. Cells were incubated for 48 hours and assayed for cell viability with Cell Titer-Glo (Promega). All well values were normalized to the average of DMSO control wells found on each plate. Figure 1A shows average normalized values of duplicate plates. Full screen results can be found in Table S1.

### Animal experiments

All animal work was approved by the Institutional Animal Care and Use Committee of the National Institute of Environmental Health Sciences.

For xenograft experiment, 2×10^6^ *M. hyorhinis*-positive or negative HCT116 human colon carcinoma cells were subcutaneously injected into each flank of 6 week old female Nu/J mice (#002019, Jackson labs). We had eight mice per treatment group and all mice developed at least one tumor. Tumor length and width were measured by caliper and tumor volume was calculated using the formula V=length*width^2^/2. Treatment with 15 mg/kg twice daily subcutaneous injections of STF118804 or vehicle control was initiated when the largest tumor reached 200 mm^3^.

Due to solubility problem of the published STF118804 formulation^22^ we developed a new formulation using block co-polymer micelles. The amphiphilic triblock copolymer [P(MeOx_35_-*b*-BuOx_20_-*b*-MeOx_34_), Mn=8.4 kg/mole, PDI =1.18)] was synthesized by living cationic ring-opening polymerization^23,24^. Briefly, pre-determined amounts of polymer and STF118804 were dissolved in acetone and mixed well at 1:10 drug:polymer ratio, followed by complete evaporation of acetone to form a thin film. The formed thin film was rehydrated with appropriate amount of distilled water and sonicated for 10 min. To remove residual solid STF118804 (if any), the samples were centrifuged at 10,000 g for 3 min and the supernatant was obtained and lyophilized. The lyophilized samples were rehydrated with normal saline immediately before use. The concentration of STF118804 in polymeric micelles was determined by reverse-phase high-performance liquid chromatography using an Agilent Technologies 1200 series HPLC system equipped with UV detector and a Nucleosil C18 5 μm column (250 mm × 4.6 mm). STF118804 was eluted with ACN/water; 70/30 as mobile phase, at a flow rate of 1mL/min and detected at 310nm. The concentration of STF118804 was calculated using a linear calibration curve against standard STF118804 solution.

Mice were sacrificed after two weeks of treatment or earlier when tumors reached 1000 mm^3^ as required by the animal protocol.

For isotopic tracing experiment, eight weeks old C57BL/6 male mice were either given regular water or autoclaved water containing antibiotic cocktail (1 g/l ampicillin, 1 g/l neomycin, 1 g/l metronidazole, 500 mg/l vancomycin) for twelve days. Fecal DNA was isolated using QIAamp DNA Stool Mini Kit (Qiagen) and microbiota depeletion was verified by qPCR using 16S V3 sequencing primers (5’-tcgtcggcagcgtcagatgtgtataagagacagccagactcctacgggaggcag-3 ‘; 5’-gtctcgtgggctcggagatgtgtataagagacagcgtattaccgcggctgctg-3’).

Control and microbiota-depleted mice were treated with oral gavage of 80 mg/kg nicotinamide labeled with 4 deuterium atoms on the pyridine ring (D4-NAM). Additional control group of mice also received a PBS gavage. Gavages were staggered and were given alternating between control and microbiota-depleted mice to avoid any experimental bias. Mice were sacrificied three hours after gavage. Colons were flushed and flash frozen in liquid nitrogen along with livers. Samples were stored at -80 degrees before processing. Each treatment group had 5-6 mice.

### Plasmids

For CRISPR-Cas9-mediated E2F1 knockout, gRNA sequence 5’-ggagatgatgacgatctgcg-3’ targeting exon 1 of E2F1 was cloned into LentiCRISPR v.2 (Addgene, #52961).

For inducible knockdown of E2F1, shRNAmir insert from E2F1-targeting pGIPZ (Dharamacon, #V3LHS_393591) was cloned into pTRIPZ vector (Dharmacon) with MluI and XhoI to produce pTRIPZ-shE2F1.

For overexpression of HA-PncA, *pncA* gene was amplified from *E.coli* genomic DNA using caccctcgaggccccctcgcgccctgttactg and caccgcggccgcttacccctgtgtctcttcccag primers and cloned into pcDNA3-HA plasmid using XhoI and NotI sites. The cloned *pncA* sequence was verified by Sanger sequencing.

### E2F1 knockdown and PncA overexpression

For E2F1 knockdown, lentiviral particles were produced using 293T cells co-transfected with pMD2.G (Addgene #12259), and psPAX2 (Addgene #12260) and LentiCRISPR v.2 targeting construct for E2F1 knockout or pTRIPZ-shE2F1 for inducible E2F1 knockdown using Mirus TransIT-LT1 Transfection Reagent. Target cells were infected with lentivirus-containing supernatant and selected with 2.5 μg/ml puromycin. Expression of the inducible shRNA from pTRIPZ-shE2F1 was induced by addition of 1 μg/ml doxycycline.

For PncA overexpression, CRC119 cells were transiently transfected with pcDNA-HA-pncA using GenJet Ver.II transfection reagent (Signagen).

### Western blot

Cell pellets were lysed with RIPA buffer containing Complete Mini protease inhibitor cocktail (Roche Diagnostics). Proteins were resolved on 4-20% gradient SDS-PAGE, transferred to PFDF membranes and probed with antibodies for E2F1 (Santa Cruz Biotechnologies, sc-251), GAPDH (Cell Signaling Technology, 2118S), alpha-tubulin (Abcam, Ab7291), HA (Santa Cruz Biotechnologies, sc-7392).

### NAD, ATP and viability enzymatic assays

For all assays, cells were plated at 10,000 cells per well in clear bottom white plates (Corning, #3610). For enzymatic measurement of total NAD, NAD/NADH-Glo™ kit (Promega) were used. CellTiter-Glo^®^ (Promega), which measures cellular ATP, was used for ATP kinetics (Fig. 2A) and as a proxy for cell number (viability).

### Metabolomics analysis using LC-MS

Cells were grown in 6-well plates in triplicates and treated with 100 nM STF118804 or DMSO control for 24 hours. After collection of medium samples, cells were briefly washed with saline and metabolites were extracted by scraping cells on dry ice into cold 80% methanol/20% water. Plates were incubated at -80 degree for 15 minutes and extracts transferred into microcentrifuge tubes. Following centrifugation at 14000 rpm for 10 minutes, supernatants were transferred to new tubes and dried in a vacuum concentrator at room temperature. The dry pellets were reconstituted into 30 μl (per 3 mg tissue) sample solvent (water:methanol:acetonitrile, 2:1:1, v/v) and 3 μl was further analyzed by liquid chromatography-mass spectrometry (LC-MS).

For tissue analysis, frozen tissues were pulverized in a mortar in liquid nitrogen and stored on dry ice. 20-30 mg of tissue powder was weighed, extracted in cold 80% methanol/20% water and processed as described for cells.

LC-MS method for metabolomics- Ultimate 3000 UHPLC (Dionex) is coupled to Q Exactive Plus-Mass spectrometer (QE-MS, Thermo Scientific) for metabolite profiling. A hydrophilic interaction chromatography method (HILIC) employing an Xbridge amide column (100 x 2.1 mm i.d., 3.5 μm; Waters) is used for polar metabolite separation. Detailed LC method was described previously^25^ except that mobile phase A was replaced with water containing 5 mM ammonium acetate (pH 6.8). The QE-MS is equipped with a HESI probe with related parameters set as below: heater temperature, 120 °C; sheath gas, 30; auxiliary gas, 10; sweep gas, 3; spray voltage, 3.0 kV for the positive mode and 2.5 kV for the negative mode; capillary temperature, 320 °C; S-lens, 55; A scan range (m/z) of 70 to 900 was used in positive mode from 1.31 to 12.5 minutes. For negative mode, a scan range of 70 to 900 was used from 1.31 to 6.6 minutes and then 100 to 1,000 from 6.61 to 12.5 minutes; resolution: 70000; automated gain control (AGC), 3 × 106 ions. Customized mass calibration was performed before data acquisition. Metabolomics data analysis-LC-MS peak extraction and integration were performed using Sieve 2.2 (Thermo Scientific). The peak area was used to represent the relative abundance of each metabolite in different samples. Missing values were handled as described in ^25^. MetaboAnalyst package was used for the PCA analysis, the differential metabolite presentation by volcano plot and for metabolic pathways enrichment analysis ^26^.

For targeted analyses of compounds from the NAD pathway, a method based upon the report from Yaku, et al. was developed ^27^. Data were acquired on a Q Exactive Plus mass spectrometer (QE-MS, Thermo Scientific) interfaced with a Vanquish (Thermo Fisher) UHPLC system. Reverse-phase chromatography was performed using a CORTECS C18 column (100 x 2.1 mm i.d., 1.6 μm; Waters) with solvent A being 5 mM ammonium formate in water (pH 6.5) and solvent B being methanol and a flow rate of 150 μL/minute. The LC gradient included a ramp from 0% to 42% B over the first 6 minutes followed by a ramp to 95% over the next minute. A 3 minute hold at 95% was followed by a return to 0% B over the next 0.5 minutes. The run was completed with a 9.5 minute recondition at 0% B. For the mass spectrometry, a PRM method was employed with a segmented include list for the masses of the metabolites of interest and their optimized normalized collision energies. The QE-MS was equipped with a HESI source used in the positive ion mode with the following instrument parameters: sheath gas, 40; auxiliary gas, 10; sweep gas, 1; spray voltage, 3.5 kV; capillary temperature, 325 °C; S-lens, 50; scan range (m/z) of 70 to 1000; 2 m/z isolation window; resolution: 17,500; automated gain control (AGC), 2 × 10e5 ions; and a maximum IT of 200 ms. Mass calibration was performed before data acquisition using the LTQ Velos Positive Ion Calibration mixture (Pierce). PRM data were processed using either the Qual Browser application or the Xcalibur processing feature in the Xcalibur software suite (Thermo Scientific). Extracted ion chromatograms for fragment ions were drawn for each of the NAD metabolites in ther respective channels at their appropriate elution times and areas under the peak calculated and used to represent the relative abundance of each metabolite in the samples. Peak areas were normalized to tissue weight.

### RNA isolation and RT-PCR

Xenograft tumors were frozen and powdered in liquid nitrogen using mortar and pestle. Total RNA was isolated using RNeasy Kit (Qiagen) and reverse-transcribed using High-Capacity cDNA Reverse Transcription Kit (ThermoFisher Scientific). Real-time PCR was performed on CFX96 real time PCR instrument (Bio-Rad) using iQ SYBR Green SuperMix (Bio-Rad) with the following primers: E2F1 (cggcgcatctatgacatcac, gtcaacccctcaagccgtc), CCNA2 (cctgcgttcaccattcatgt, cagggcatcttcacgctctat), GAPDH (acccactcctccacctttga, ctgttgctgtagccaaattcgt).

### Mitochondrial membrane potential loss

Cells were trypsinized, stained with 100 nM tetramethylrhodamine ethyl ester (TMRE) and analyzed by flow cytometry on LCR II instrument (BD Biosciences).

### NAD measurement in isolated cytosolic and mitochondrial fractions

Mitochondrial and cytosolic fraction were isolated according to^28^ and relative NAD/NADH content was determined with NAD/NADH Quantitation Kit (Sigma, MAK037). NAD values were normalized to protein content determined by BCA kit (ThermoFisher Scientific).

### Bioenergetic profiling

Extracellular acidification rate (ECAR) and oxygen consumption rate (OCR) were analyzed on Seahorse XFe96 Analyzer (Agilent) using Seahorse XF Cell Mito Stress Test Kit and Glycolysis Stress Test Kit (Agilent). Following completion of the analysis, protein content in each well was determined by BCA kit (ThermoFisher Scientific) and used for normalization of the raw ECAR and OCR values.

