## Supplementary material for "Bacteria boost mammalian host NAD metabolism by engaging the deamidated biosynthesis pathway"

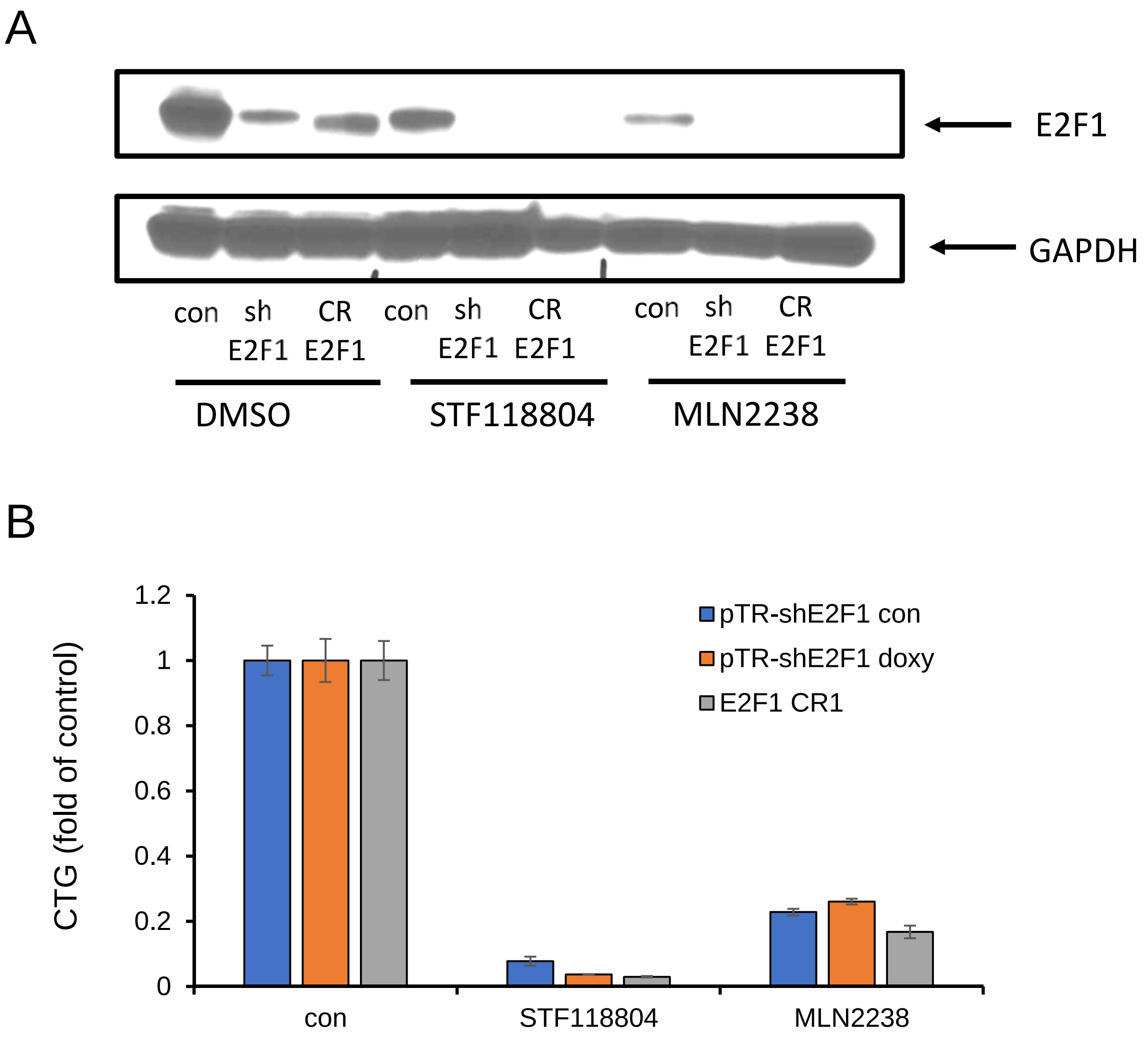


**Figure S1. E2F1 knockdown does not affect sensitivity to STF118804, MLN2238.** E2F1 was knocked down in CRC240 cells either using a doxycycline-inducible shRNA (doxy) or CRISPR/cas9 genome editing (CR). Cells were treated with 100 nM STF118804 or 100 nM MLN2238 for 41 hours. E2F1 protein levels were analyzed by Western blot (A) and cell viability was measured by CTG assay (B). Data are means and SD of biological triplicates from a representative experiment repeated twice with similar results.

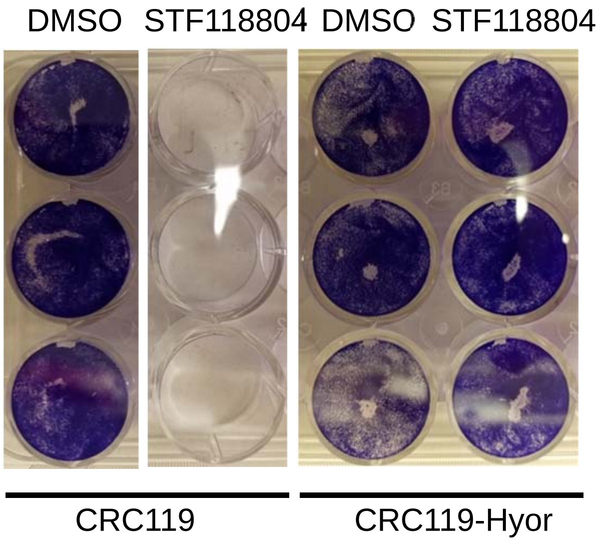


**Fig. S2. *Mycoplasma hyorhinis* confers mammalian cells with resistance to STF118804.** CRC119 cells or CRC119-Hyor cells chronically infected with Mycoplasma *hyorhinis* were seeded in 12-well plates. Next day, cells were treated with 100 nM STF-118804. Seventy-two hours later cells were washed and adherent cells were stained with crystal violet.

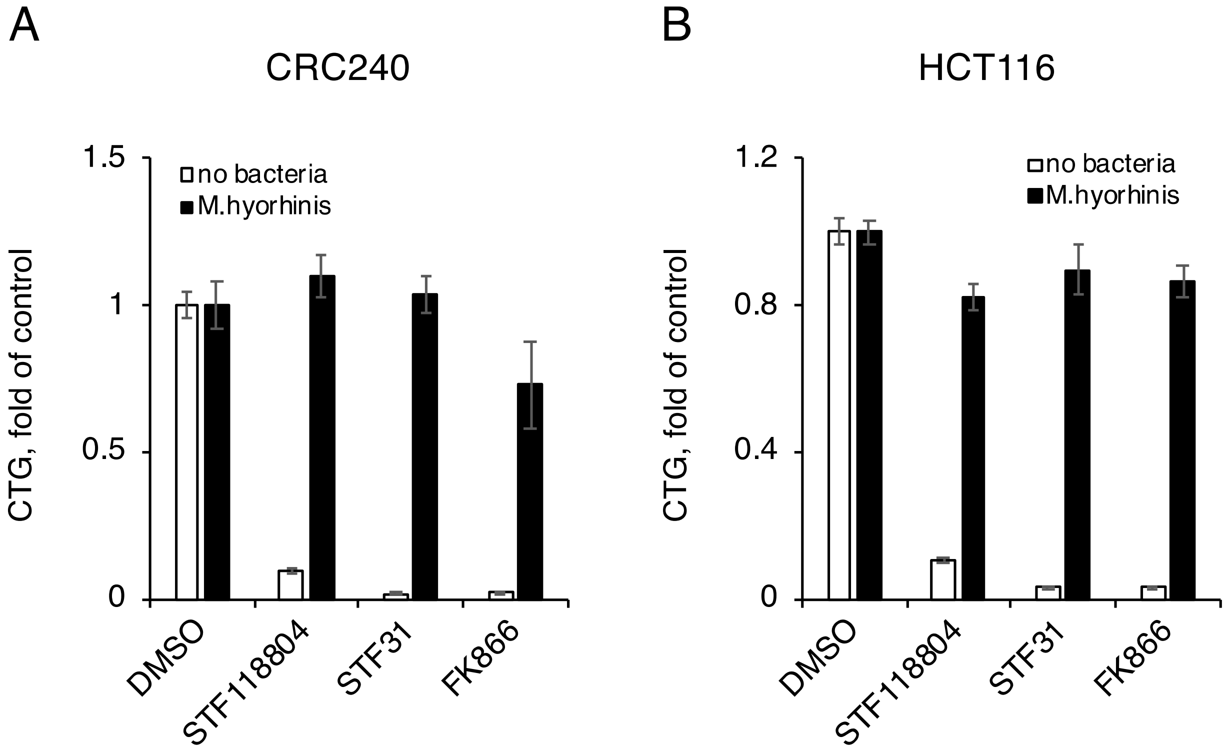


**Figure S3. Mycoplasma protects from NAMPT inhibitors in additional cell lines.** CRC240 and HCT116 colon cancer cell lines either without bacteria or chronically infected with *M.hyorhinis* were treated with three different NAMPT inhibitors (100 nM STF118804, 1 μM STF31 or 40 nM FK866) for 48 hr. Cell viability was measured by CTG assay. Data are means and SD of biological triplicates from a representative experiment repeated twice with similar results.

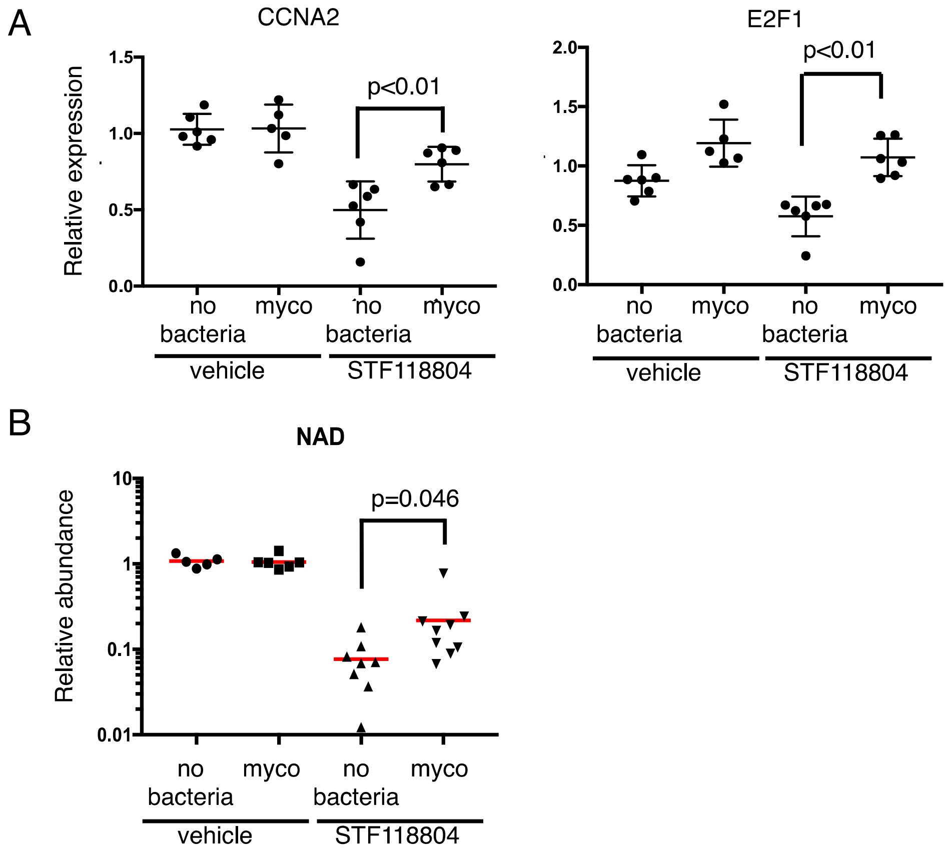


**Figure S4. Mycoplasma protects xenograft tumors from NAMPT inhibitors induced repression of proliferation genes and NAD depletion in mice.** (A) Mycoplasma prevents repression of proliferation genes by a NAMPT inhibitor. Clean (no bacteria) or *M.hyorhinis*-infected HCT116 cells were xenografted into nude mice. Mice were treated with STF118804 or vehicle control as described in Materials and Methods. The mRNA levels of Cyclin A2 (CCNA2) and E2F1 in tumors were quantified by qPCR. The p-value was calculated by two-tailed Mann-Whitney U-test (n=5-6).
(B) Mycoplasma attenuates NAD depletion in tumors of STF118804-treated mice. Tumor NAD levels were quantified by LC-MS and normalized to kynurenine. The p-value was calculated by one-tailed t-test (n=8-9).

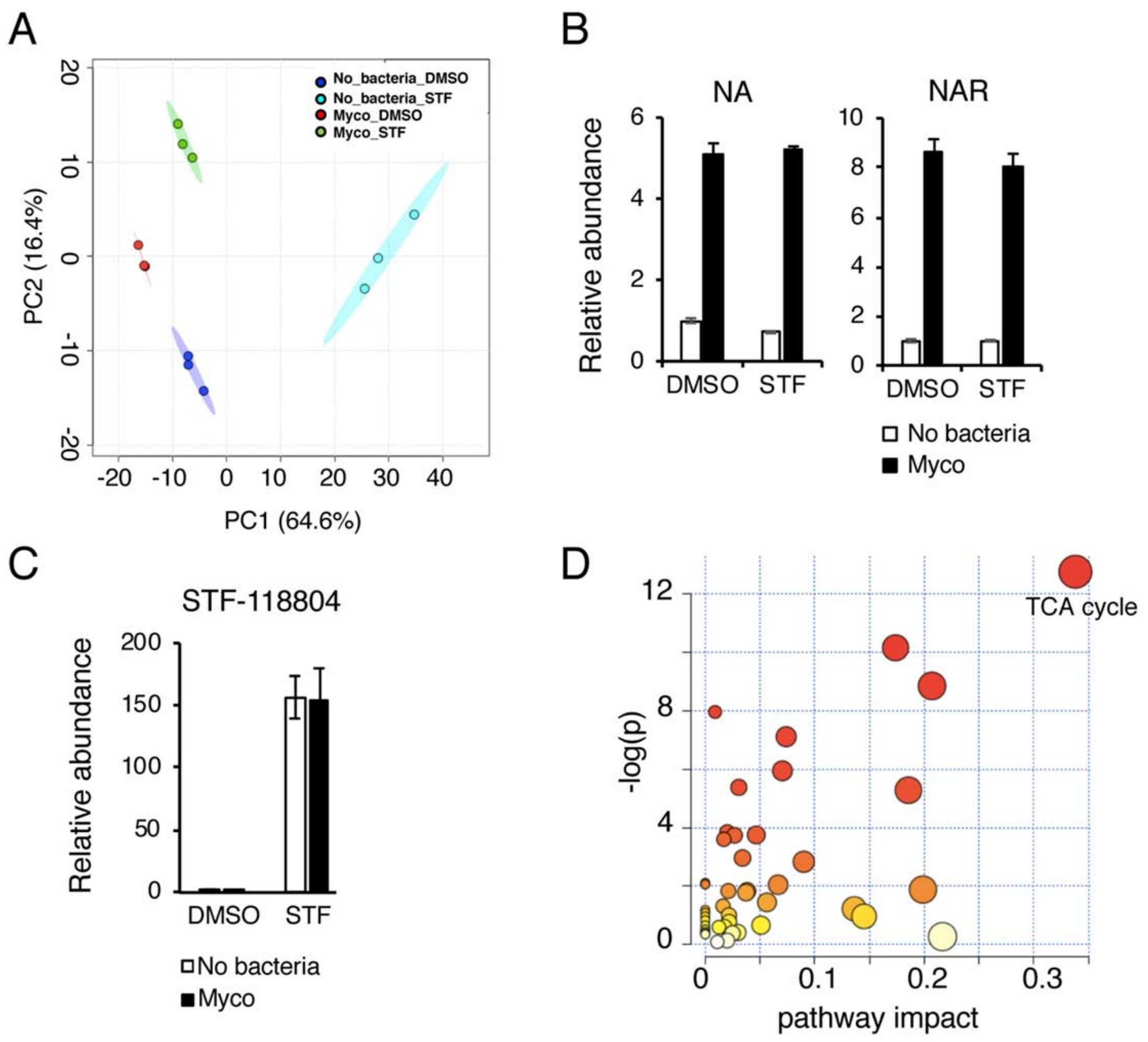


**Figure S5. Metabolomics of CRC119 infected with *M. hyorhinis.*** (A) Principal component analysis. (B) Mycoplasma increase the concentration of Nicotinic acid (NA) and Nicotinic acid riboside (NAR) in the culture medium. Metabolites were analyzed by LC-MS. (C) Mycoplasma do not affect the intracellular levels of STF118804. The concentrations of intracellular STF-118804 were analyzed by LC-MS. (D) The TCA cycle is the most significantly altered metabolic pathway induced by mycoplasma in STF118804-treated cells. Bar graphs represent means and standard deviations of triplicate samples from a single experiment.

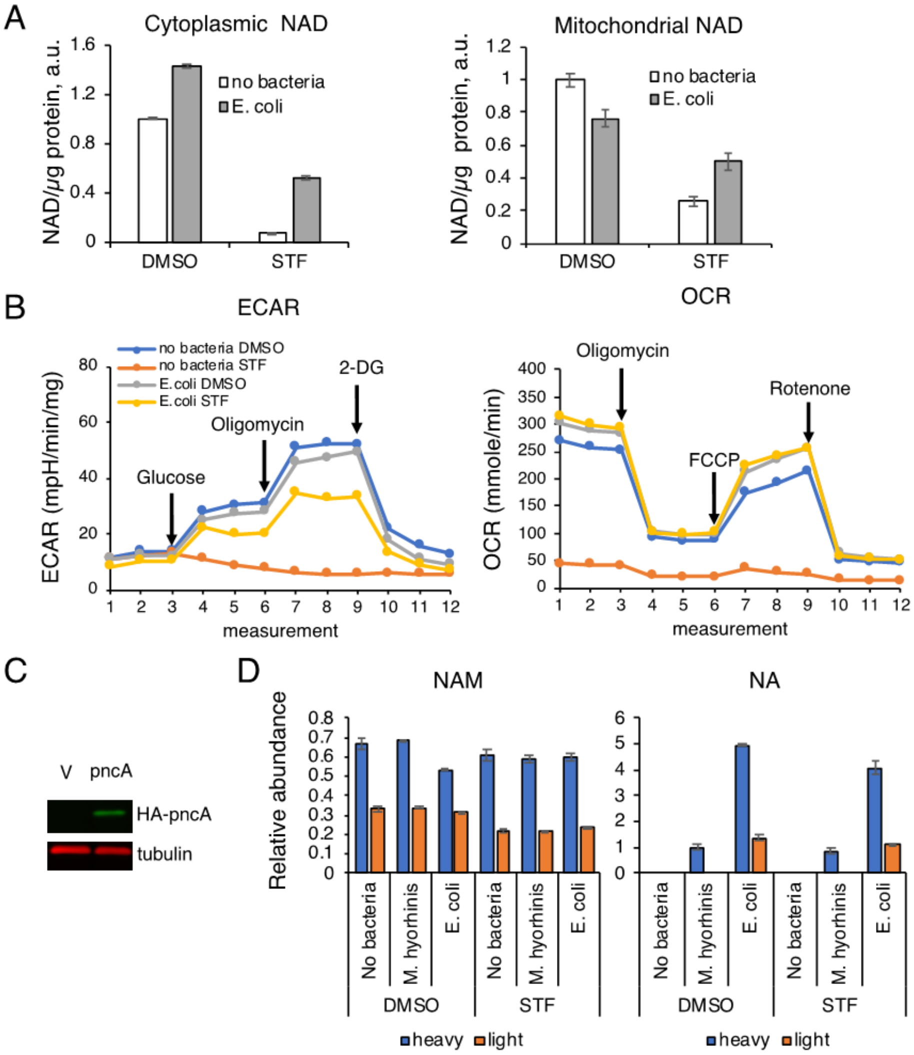


**Figure S6. E. coli rescue NAMPTi-induced toxicity through pncA**. (A) E. coli rescue STF118804-mediated depletion of cytoplasmic and mitochondrial NAD. CRC119 cells were treated with 100 nM STF18804 or DMSO control with or without 1:1000 dilution of overnight E. coli culture in the presence of 1 µg/ml gentamycin for 22 hours. Relative levels of total NAD (NADH+ NAD+) were measured in isolated cytoplasmic and mitochondrial fractions by a colorimetric enzymatic assay and normalized to total protein. **(B)** E. coli rescue STF118804-mediated shutdown of energy metabolism. CRC119 cells were infected with 1:1000 dilution of overnight E. coli culture (with 1 µg/ml gentamycin) or with control medium and cells were treated with 100 nM STF118804 or DMSO control. Seahorse analysis using Glycolysis Stress Test or Mito Stress Test was performed 40 hr later. Raw ECAR readings were normalized to total protein content. (C) Overexpression of E. coli pncA in CRC119 cells. Western blot of HA-pncA and tubulin loading control. (D) Bacteria enhances conversion of NAM into NA. CRC119 cells were infected with the indicated bacteria and treated with 200 nM STF118804 or DMSO control for 24 hours in the presence of 5 mg/l D4-labeled NAM. The relative abundance of unlabeled (light) and labeled (heavy) NA and NAM was measured by LC-MS.
Results are means and standard deviations of triplicate samples from representative experiments (N=3-6).

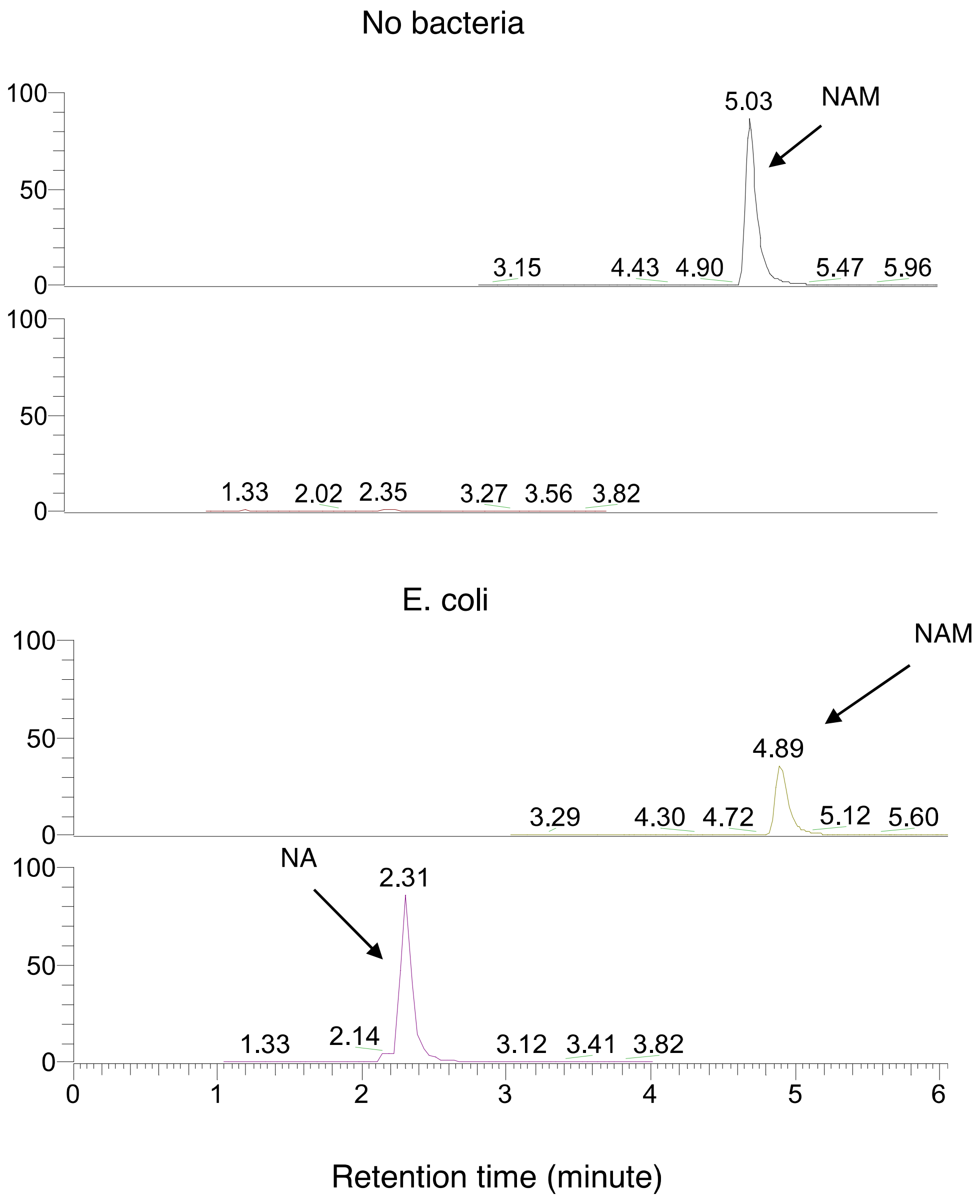


**Figure S7. E. coli induce the conversion of NAM to NA in culture medium.**
Representative chromatograms from the LC-MS analysis of NAM-containing RPMI medium after two hours of incubation at 37 degrees with or without *E. coli*. Bacteria were removed by centrifugation and filtering prior to the analysis.

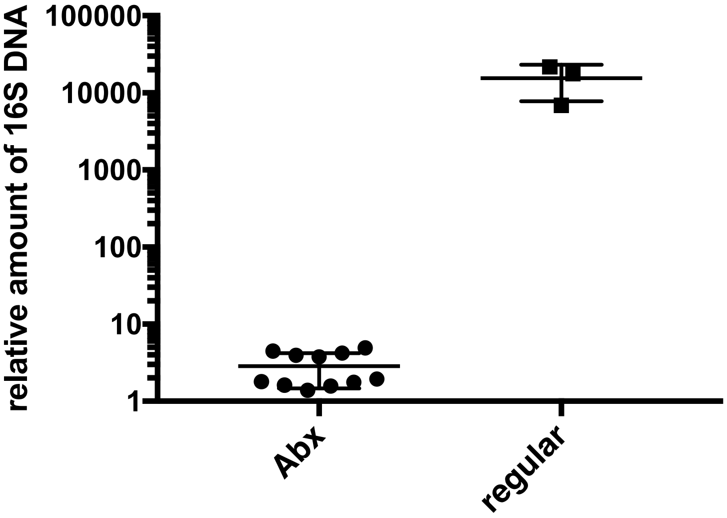


**Figure S8. Depletion of gut microbiota in mice by antibiotic treatment.** qPCR analysis of 16S rDNA in fecal DNA of mice used in the experiment described in Fig. 4H and 4I. All microbiota-depleted mice and three representative conventional mice were analyzed.
